# Generation of tonsil organoids as an *ex vivo* model for SARS-CoV-2 infection

**DOI:** 10.1101/2020.08.06.239574

**Authors:** Han Kyung Kim, Hyeryeon Kim, Myoung Kyu Lee, Woo Hee Choi, Yejin Jang, Jin Soo Shin, Jun-Yeol Park, Seong-In Hyun, Kang Hyun Kim, Hyun Wook Han, Meehyein Kim, Young Chang Lim, Jongman Yoo

**Affiliations:** Department of Microbiology, CHA University School of Medicine, Seongnam, Republic of Korea; CHA Organoid Research Center, CHA University, Seongnam, Republic of Korea; R&D Institute, ORGANOIDSCIENCES LTD, Seongnam, Republic of Korea; Department of Otorhinolaryngology-Head and Neck Surgery, the Research Institute, Konkuk University School of Medicine, Seoul, Republic of Korea; Infectious Diseases Therapeutic Research Center, Korea Research Institute of Chemical Technology, Daejeon, Republic of Korea; Department of Biomedical informatics, CHA University School of Medicine, CHA University, Seongnam, Republic of Korea; Graduate School of New Drug Discovery and Development, Chungnam National University, Daejeon, Republic of Korea

## Abstract

Palatine tonsil (hereinafter referred to as “tonsil”) plays role in the immune system’s first line of defense against foreign pathogens. Coronavirus disease 2019 (COVID-19), caused by severe acute respiratory syndrome coronavirus 2 (SARS-CoV-2), has become a worldwide pandemic since the infection was first reported in China in December 2019. The aim of this study was to establish tonsil epithelial cell-derived organoids and to examine their feasibility as an *ex vivo* model for SARS-CoV-2 infection. Using an optimized protocol, we achieved 3D tonsil organoid culture from human tonsil tissue that reflects the distinctive characteristics of the tonsil epithelium, such as its cellular composition, histologic properties, and molecular biological features. Notably, we verified that SARS-CoV-2 can infect tonsil organoids with a robust replication efficiency. Furthermore, treatment with remdesivir, an antiviral agent, effectively protected them from viral infection. Therefore, tonsil organoids could be available for investigation of SARS-CoV-2 infection-mediated pathology and for preclinical screening of novel antiviral drug candidates.

**One-sentence Summary:** This study established tonsil epithelial cell-derived organoids and demonstrated their feasibility as an *ex vivo* model for SARS-CoV-2 infection.

## Introduction

COVID-19 is rapidly spreading worldwide, resulting in over 14 million infected cases with approximately 597 thousand death as of late July 2020 (https://www.who.int/docs/default-source/coronaviruse/situation-reports/20200719-covid-19-sitrep-181.pdf?sfvrsn=82352496_2)(2020). SARS-CoV-2 is mainly transmitted through the upper respiratory tract with diverse disease symptoms ranging from mild sickness (including fever, cough, sore throat, and loss of olfaction or taste) to radical severity (pneumonia and even death)(Delgado-Roche and Mesta, 2020; Huang et al., 2020; Wang et al., 2020; Xu et al., 2020). Thus, development of preventive vaccines or therapeutic agents against SARS-CoV-2 is urgently needed. Regarding this, a suitable *ex vivo* model representing human infection cases in terms of host immune responses as well as antiviral pathways is indispensable for preclinical validation of efficacy and toxicity.

The tonsils are two oval-shaped immunologic organs in the back of the oral cavity(Perry and Whyte, 1998) (Figure S1A). They are histologically composed of mucosa-lining epithelium and underlying parenchymal B or T cell lymphocytes(Perry and Whyte, 1998). These cells filter diverse inhaled viruses, possibly including SARS-CoV-2(Nave et al., 2001; Perry and Whyte, 1998). Thus, it was assumed that the tonsil could be one of the most suitable and easily accessible organs for studying respiratory virus-mediated diseases, such as SARS-CoV-2 infections. Currently, there are no substitutable model systems for exploring tonsillar infectious diseases because of the difficulty in culturing its epithelial cells that can be stably maintained over long-term serial passages(Kang et al., 2015). In addition, experimental small animal models are inappropriate due to intrinsic lack of tonsils in mice or hamsters(Casteleyn et al., 2011).

The tonsil epithelium is composed of differentiated epithelial cells, which reside in the top layer of the epithelium and are generated by the differentiation of stem-like epithelial cells in the bottom layer of the epithelium(Kang et al., 2015). This suggests that producing an organoid model could be feasible, particularly given recent advances in long-term 3D culture of epithelial stem cells from human specimens(Huch et al., 2013; Sato et al., 2009). These epithelial organoids mimic the *in vivo* environment because they contain all tissue-specific epithelial cell types and structures that differentiate and proliferate from epithelial stem cells(Bartfeld et al., 2015; Clevers, 2016; Dutta et al., 2017; Huch et al., 2015; Ren et al., 2014; Sato et al., 2011; Zhang et al., 2020). This method has been extensively used to generate epithelial tissue-related disease models and study epithelial biology(Bartfeld et al., 2015; Clevers, 2016; Lancaster and Knoblich, 2014; van de Wetering et al., 2015).

Here, we aimed to establish tonsil-derived epithelial organoids with morphological and molecular biological features similar to those of *in vivo* tonsils and to examine their availability as an *ex vivo* model for SARS-CoV-2 infection.

## Results

### Generation of human tonsil organoids from tonsil tissues

To generate epithelial organoids from human tonsils, enzymatically dissociated cells from human tonsil tissue were embedded in Matrigel and grown in either human colon organoid medium (HCM)(Drost et al., 2015) or human prostate organoid medium (HPM)(Karthaus et al., 2014) containing epidermal growth factors, such as Noggin, fibroblast growth factor (FGF) 2, FGF10, and R-spondin-1, which play pivotal roles in differentiation and proliferation of epithelial stem cells (Figure S1A and Table S1). Tonsil cells cultured in HPM yielded a higher number of 3D epithelial organoids with a stratified epithelial structure within 10 days, compared to the cells cultured in HCM (Figure S1B, C). To optimize the medium composition, we screened the medium components. The largest organoids in the highest formation rate were observed when dissociated tonsil cells were grown in the HPM in the absence of epidermal growth factor, defined as the tonsil organoid formation medium (TfM) (Figure 1A, B and S1D-F). However, we found that subculture of the tonsil organoids in TfM was limited within two passages and further expansion was not available (Figure S1G). To solve this problem, we additionally modified the TfM in which the p38 pathway is activated by removing SB202190 and adding hepatocyte growth factor (HGF), defined as the human tonsil expanding medium (TeM). In this medium, the organoids were successfully expanded beyond 60 days without affecting the growth rate (Figure 1C, D and S1H, I).

**Figure 1.**
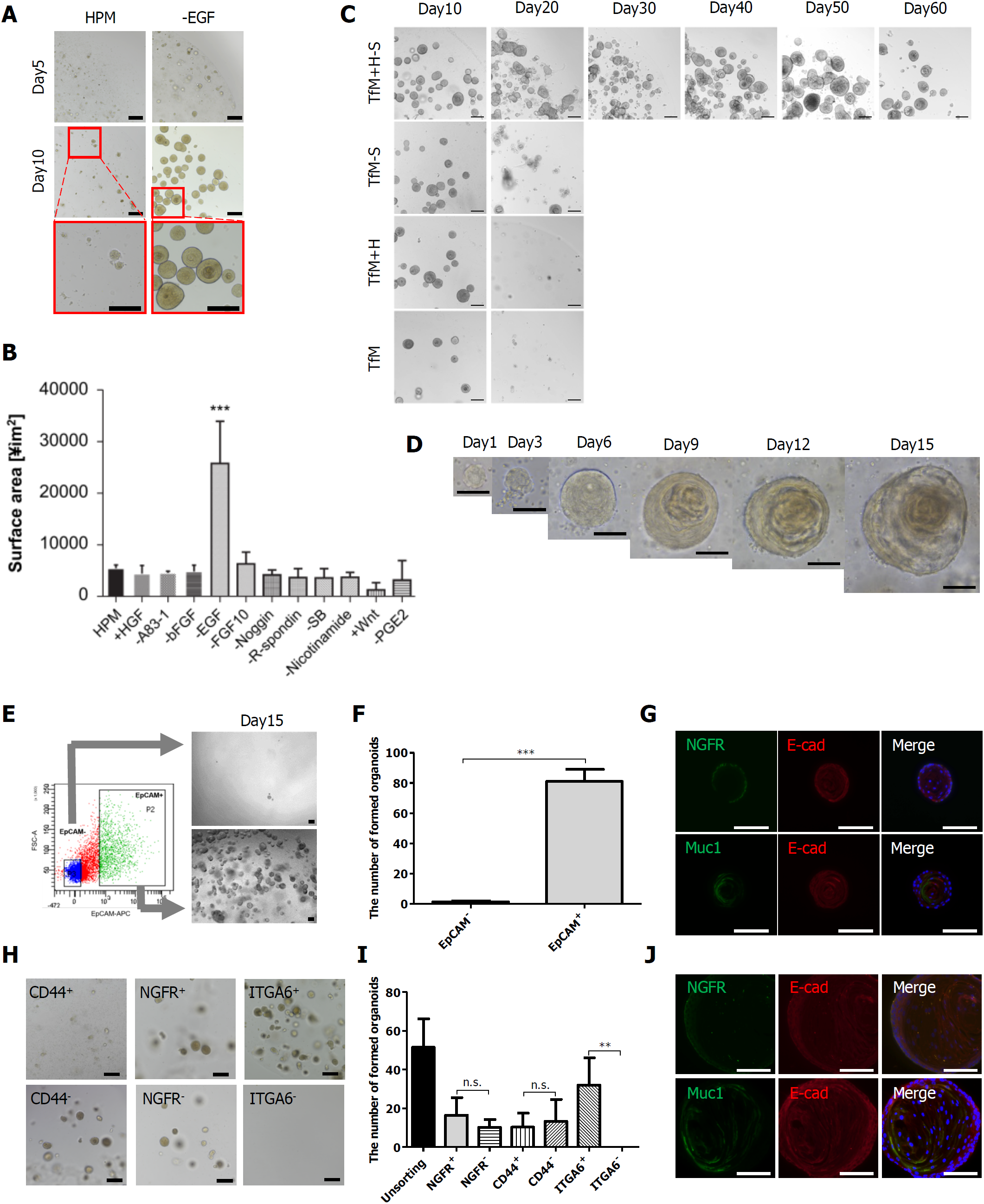
Generation of human tonsil organoids from tonsil tissues. (A) Comparison of tonsil organoids formation in human prostate media (HPM) with and without EGF. (B) The size of organoids cultured in HPM containing different combinations of factors. Values are expressed as mean ± SEM of three independent experiments. *** means p < 0.001. (C) Representative images of human tonsil organoids cultured in TfM (tonsil organoid formation medium), TfM with HGF (abbreviated to +H), TfM without SB202190 (-S), and TfM with HGF and without SB202190 (+H-S). (D) Time-dependent morphology and growth of tonsil organoid cultured in TeM (tonsil organoid expansion medium, TfM+H-S). (E-G) Formation and characterization of tonsil organoid in the FACS-sorted EpCAM^+^ or EpCAM^-^ cells population (E,F). *** means p < 0.001. Immunofluorescence images indicate the expression of tonsil specific markers such as nerve growth factor receptor marker (NGFR), cell surface marker (Muc1), and epithelial marker (E-cadherin) in FACS-purified EpCAM^+^ cells in TeM from day 15 (G). (H-J) Different organoid formation in the FACS-sorted CD44^+^, NGFR^+^, or ITGA6^+^ cells (H,I), and expression of tonsil specific markers in the ITGA6+ derived organoids (J). ** means p < 0.01. The number of formed organoids represents the mean ± SEM of the three independent experiments. In all panels, scale bar, 100 µm.

Epithelial cell adhesion molecule (EpCAM), a broad epithelial cell marker(Trzpis et al., 2007), is expressed in the basal layer of the tonsil epithelium and has been reported to exist on the outer surface of other organoids(Buzhor et al., 2011; Itoh, 2016; Jung et al., 2011; Kessler et al., 2015). We examined whether the tonsil organoids originated from EpCAM^+^ cells by sorting cells for culture. In contrast to EpCAM^-^ cells, EpCAM^+^ ones efficiently formed organoids (Figure 1E, F). These EpCAM^+^ organoids substantially exhibited structural similarity to the tonsil organoids grown from unsorted cells as assessed by immunofluorescence staining for NGFR, MUC1, and E-cadherin (Figure 1G) The result indicated that the majority of tonsil organoids are differentiated from EpCAM^+^ epithelial cells.

To investigate which tonsil markers among ITGA6, NGFR, and CD44 are critical for organoid maturation, cells were sorted again on the basis of expression level of the individual markers(Kang et al., 2015)(Figure S2A). The efficiency of organoid formation was highest for ITGA6^+^ cells, while no organoid formation was observed with ITGA6^-^ cells (Figure 1H, I). Organoids were successfully developed irrespective of NGFR or CD expression. Expectedly, organoids derived from ITGA6^+^ cells reproduced the structural characteristics of the tonsil epithelium (Figure 1J). These findings suggest that tonsil organoids are differentiated from EpCAM^+^ and ITGA6^+^ cells.

### The tonsil organoids recapitulate features of human tonsil tissues

Next, to determine whether the tonsil organoids recapitulated the skeletal properties as well as architecture of biomarkers identifying the tonsil epithelium, cultured tonsil organoids and human tonsil tissues were compared by histological analysis (Figure 2A,B and S2B). Immunohistochemistry showed that maturation of the organoids is accomplished when the highly ordered stratified epithelium constructed for 15 days as observed in the tonsil epithelium (Figure 2B and S2B). These results were more clarified by immunofluorescence staining with specific markers representing each layer. NGFR-, CD44-, and CK5-positive cells, located in the basal layer of tonsil tissues, were observed on the outer side of organoids; MUC1- and CK4-positive cells, in the mid-upper layer of tonsil tissues, were observed on the inner side of organoids (Figure 2C and S2C). This suggests that the outer layer of the tonsil organoid was similar to the basal layer of tonsil tissue, while its inner layer is composed of cells similar to the upper layer of tonsil tissue. The proliferating cell marker Ki67 and the tonsil stem cell marker NGFR were found at the outermost layer of the organoid, and BrdU-labeled cells migrated towards the center of the organoids, confirming the inward direction of differentiation (Figure 2D and S2D). Electron Microscopic images of the organoids revealed typical features of stratified squamous epithelium and tonsil-specific elements, such as desmosomes, keratohyalin granules, and tonofilaments(Howie, 1980) (Figure 2E). Analysis of the global gene expression profiles of tonsil organoids at day 15 from 2 donors with a reference of tonsil epithelium revealed that 14,483 genes (79.1%) were similarly expressed, but 3826 genes (20.9%) were discriminatively expressed either in the tonsil epithelium (GEO accession no. GSM69767)(Kang et al., 2015) or in the tonsil organoids (Figure 2F,G).

**Figure 2.**
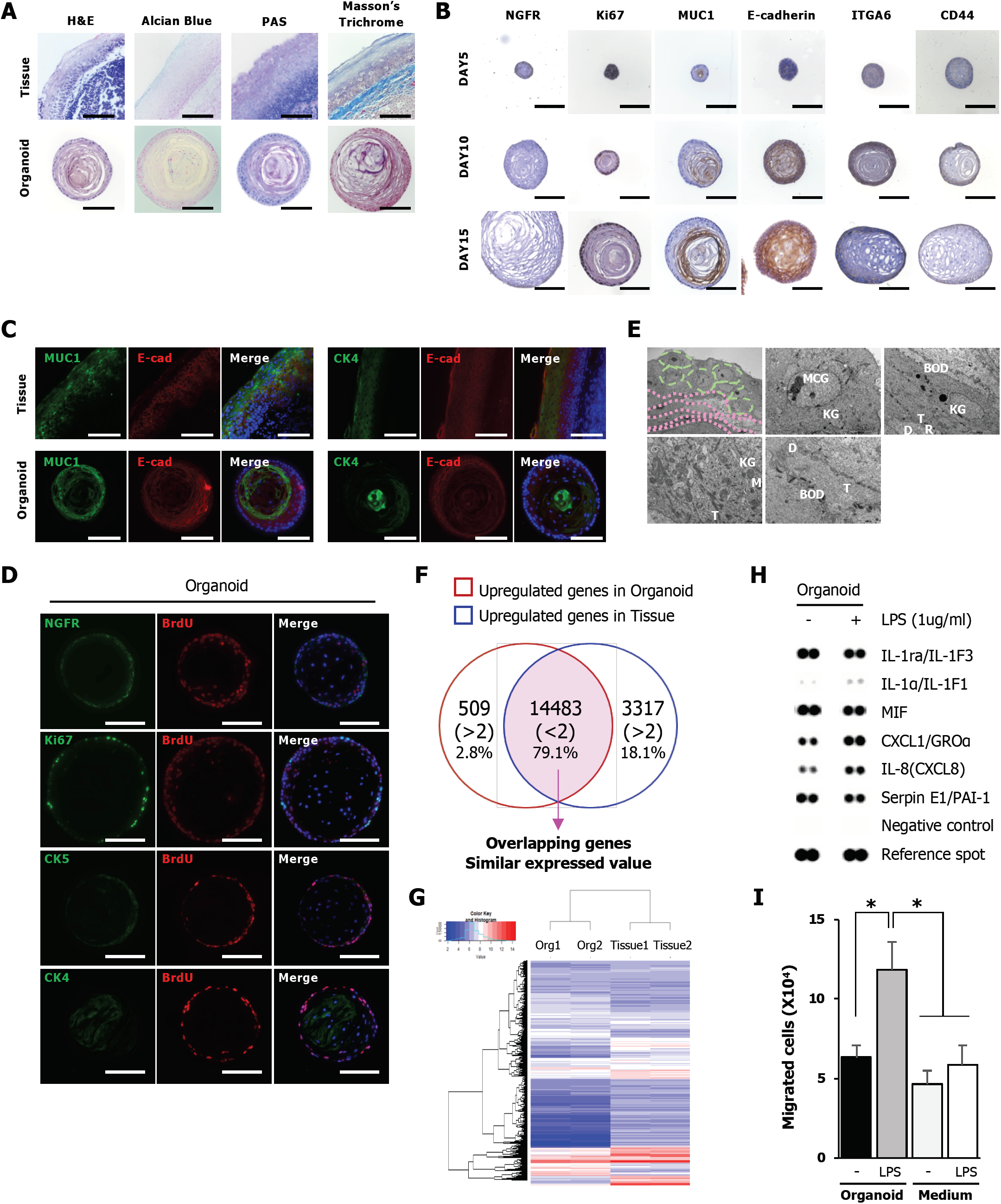
The tonsil organoids recapitulate features of human tonsil tissues. (A) Histological analyses of human tonsil epithelium and organoid using H&E staining for morphological analysis, Alcian blue staining for the mucosal layer and mucous-producing goblet cells, PAS staining for differentiated cells and Masson’s trichrome staining for connective matrix. (B) Immunohistochemistry of tonsil organoids on days 5, 10, and 15 with tonsil epithelium markers (CD44, NGFR, E-cadherin, Muc1, ITGA6) and a proliferation marker (Ki-67). (C) Immunofluorescence of tonsil epithelium and organoids cultured for 15 days in TeM with a MUC1 as mid-upper layer marker, CK4 as upper layer marker, and E-cadherin as pan-epithelial marker. (D) Immunofluorescence staining of NGFR, Ki67, cytokeratin markers (CK4, CK5), and proliferating cell marker (BrdU) in organoids treated with BrdU for two days from day 10. (E) Images of transmission electron microscopy illustrating ultrastructural morphology in human tonsil organoids. MCG: Membrane-Coating Granules, D: Desmosomes, KG: Keratohyaline Granule, R: Ribosomes, M: Mitochondria, TF: Tonofilament, T: Tonofibril. (F and G) Venn diagram (F) and Heatmap (G) showing the gene expression overlap and differentially expressed genes, respectively, between tonsil epithelium (GSE69767) and organoids (p-value < 0.05, fold change ≥ |2.0|). (H and I) Human cytokine array (H) and HL-60 migration assay (I) using LPS-stimulated tonsil organoid conditioned medium. Values are presented as the mean ± SEM of the three independent experiments. * means p < 0.05. In all panels, scale bar, 100 µm.

Tonsil epithelial cells are stimulated by microbial challenge to secrete CXC chemokines such as CXCL1, which mobilize inflammatory cells(Sachse et al., 2005), contributing to the immune system’s first line of defense against pathogens. To confirm that these responses occur in tonsil organoids, we quantified the amount of secretory cytokines after treatment with lipopolysaccharide (LPS), a cell wall component from gram-negative bacteria that causes strong immune responses. In LPS-challenged organoids, not only the expression of chemokine ligand 1 (CXCL1) and interleukin-8 (IL-8) increased at the mRNA level in cell lysates but also secretion of their proteins was enhanced in the culture supernatant (Figure 2H and S3A). The migration index of HL-60 cells was also upregulated significantly compared with the negative control following treatment with conditioned medium derived from LPS-treated organoids, indicating that LPS treatment of the tonsil organoids stimulated the secretion of chemo-attractants (Figure 2I and S3B). These results are consistent with those obtained in tissue samples from patients with tonsillitis(Sachse et al., 2005), suggesting that the tonsil organoids recapitulate tonsil epithelial responses to infectious stimuli from bacteria.

### Sustainable infection of human tonsil organoids with SARS-CoV-2

It has been reported that for attachment and infection of SARS-CoV-2, the expression of ACE2 and TMPRSS2 in the host cells is essential(Hoffmann et al., 2020; Kuhn et al., 2004). Thus, we explored whether they are expressed in tonsil organoids. Immunofluorescence analysis of ACE2 and TMPRSS2 visualized that their expression was highest in NGFR-positive basal cells and it decreased as the cells differentiated into CK5- and Muc1-positive cells (Figure 3A,B). Interestingly, in contrast to ACE2, TMPRSS2 mRNA expression increased as the tonsil organoid matured (Figure 3C,D). Collectively, we assumed that the tonsil organoids could be susceptible to SARS-CoV-2 infection.

**Figure 3.**
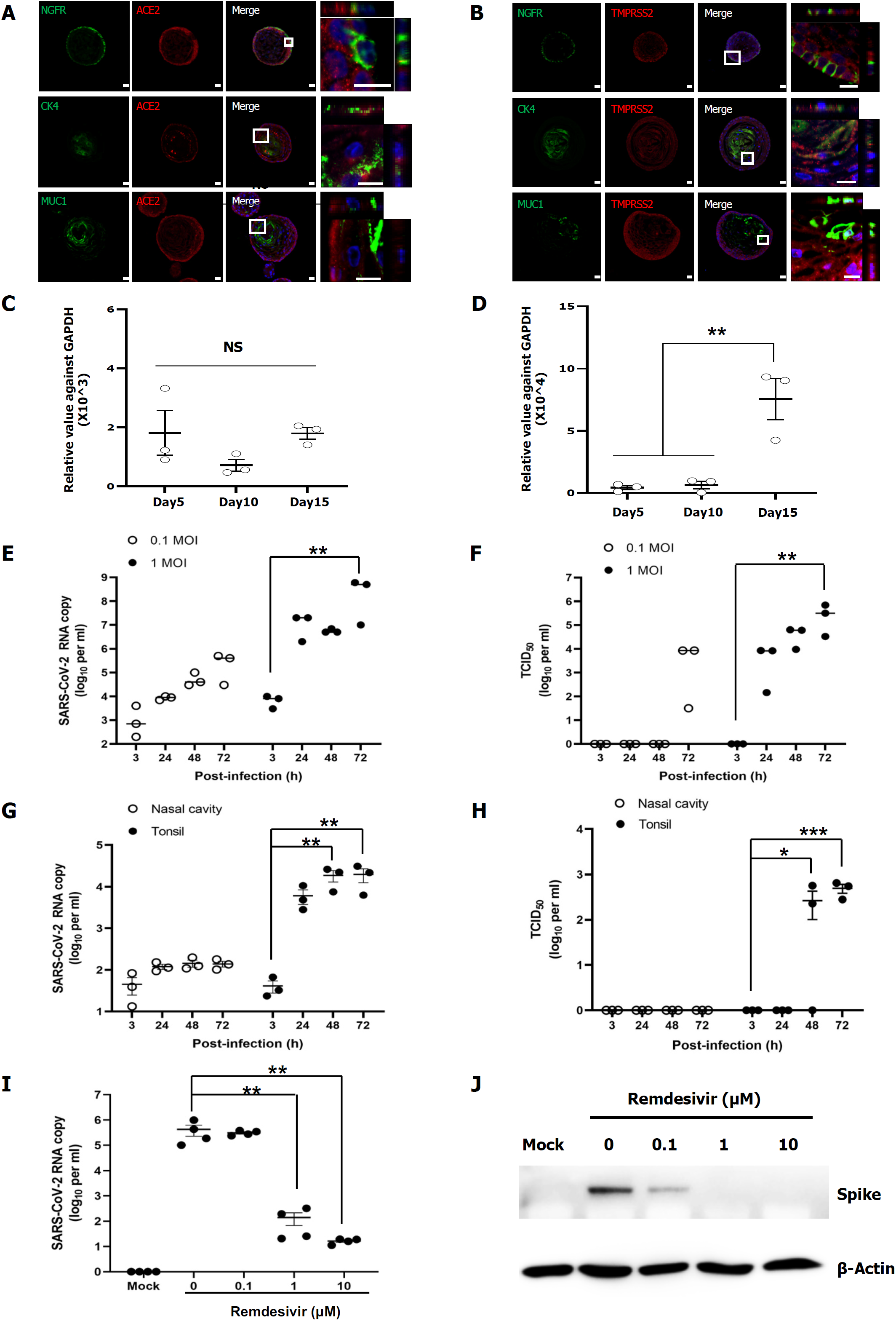
Sustainable infection of human tonsil organoids with SARS-CoV-2. (A) Immunofluorescence staining of ACE2 with NGFR as a basal layer marker, CK4 as an upper layer marker, or MUC1 as a mid-upper layer marker in tonsil organoids. (B) Immunofluorescence staining of TMPRSS2 with NGFR, CK4, or MUC1. All scale bar, 10 µm. (C and D) Quantitative RT-PCR for detecting ACE2 (C) and TMPRSS2 (D) expression in tonsil organoids originating from three different donors on days 5, 10 and 15. **, *P* < 0.01. (E) Quantitative RT-PCR for detecting viral RNA in the culture supernatants of SARS-CoV-2-infected tonsil organoids at an MOI of 0.1 or 1 at different time points after infection. **, *P* < 0.01. (F) Determination of the amount of infectious viral particle in SARS-CoV-2-infected tonsil organoids from the same donor of (E). **, *P* < 0.01. (G) Comparison of viral RNA amount existing culture supernatants from SARS-CoV-2-infected tonsil organoids and nasal cavity organoids by quantitative RT-PCR. (H) Determination of TCID_50_ of the same samples used in (G). *, P < 0.05.,***, *P* < 0.001. (I) Quantitative RT-PCR using culture supernatants of SARS-CoV-2-infected on day 2 after treatment remedsivir. Mock, no infected control. **, *P* < 0.01. (J) Western blot analysis using cell lysates to probe SARS-CoV-2 spike protein after treatment of increasing concentrations of redesivir. β-Actin was used as a loading control.

To prove the hypothesis above, the tonsil organoids were challenged with SARS-CoV-2 at different titers (0.1 and 1 MOI), and viral amplification was assessed by quantitative RT-PCR and TCID_50_ determination using the culture supernatants. RT-PCR exhibited a time-dependent increase in viral RNA levels in both challenges, indicating that the tonsil organoids are infected with SARS-CoV-2 and then release viral progeny through multi-round infections (Figure 3E). However, viral RNA detection may have occurred because of infected-organoid lysate contamination. Thus, the number of viral particles was further determined by infection of the same supernatants into fresh Vero cells. Marked enhancement of viral titers at 72 h after the lower titer challenge supported the existence of infectious SARS-CoV-2 particles in the culture supernatant (Figure 3F). Decisively, the higher titer challenge resulted in abundant accumulation of progeny virus at 24, 48, and 72 h increasingly. Next, based on the fact that the sample collection for the COVID-19 diagnostic test is generally performed through the nasal or oral mucosa swab, we compared the SARS-CoV-2 replication capacity in tonsil organoids to that in nasal cavity mucosa-derived organoids from a same donor (Figure S4A,B). The data from both RT-PCR and TCID_50_ determination showed that SARS-CoV-2 infects tonsil organoids but not nasal cavity organoids (Figure 3G,H). Taken together it can be suggested that SARS-CoV-2 infection and amplification occurs selectively in the tonsil organoids rather than in the nasal cavity organoids. Finally, we examined whether the tonsil organoid model is applicable to evaluation of antiviral activity against SARS-CoV-2 infection. The virus-infected organoids were treated with increasing concentrations of remdesivir, of which emergency use has recently been authorized by the FDA for COVID-19 treatment. Notably, a dose-dependent decrease in the viral RNA level was induced by remdesivir with remarkable inhibitory efficacy at concentrations above 1 μM (Figure 3I). Supporting this finding, western blot analysis with cell lysates from the cognate samples revealed inhibition of spike protein expression without affecting host protein expression (Figure 3J). Collectively, tonsil organoids could be applied to evaluation of the antiviral efficacy of small molecules before attempting their translation to clinical trials, as the organoid reflects the physiological conditions of viral infection in humans.

### Release of progeny SARS-CoV-2 particles and virus-induced gene expression profiles in human tonsil organoids

To define which part of the tonsil organoids is targeted by SARS-CoV-2, we immunostained the virus-infected tonsil organoids. The viral spike and dsRNA, a marker for RNA-dependent RNA replication, were dominantly expressed in the basal layer of the tonsil organoids, whereas marginal or little expression was observed the inner part of the organoids (Figure 4A,B). To further verify whether this infection indeed can cause generation of progeny viral particles, transmission electron microscopy was performed with SARS-CoV-2-infected organoids at 48 h post-infection. A high density of viral particles was observed at the outside side of the apical membrane in basal layer cells of the infected-organoids but not in the mock-infected sample (Figure 4C). These results strongly demonstrate that replication of SARS-CoV-2 in tonsil organoids facilitates generation of progeny virions.

**Figure 4.**
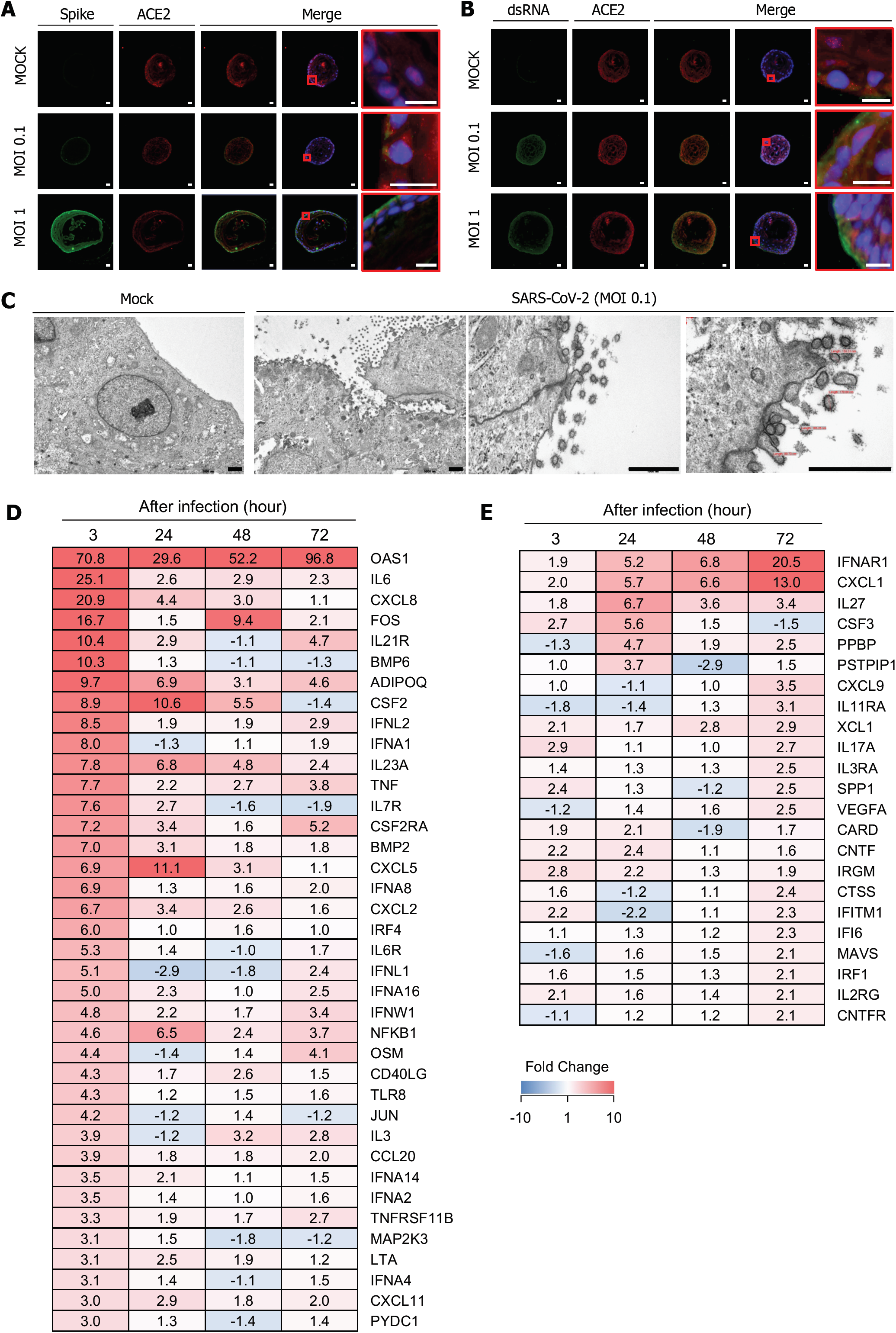
Release of progeny SARS-CoV-2 particles and virus-induced gene expression profiles in human tonsil organoids. (A and B) Immunofluorescence staining for viral spike protein (A) and viral dsRNA (B) with ACE2 in SARS-CoV-2-infected tonsil organoids on day 3 post-infection. Scale bar, 10 µm. The selected areas highlighted red boxes are also presented in merged, zoomed-in images. Nuclei were counterstained with DAPI. (C) TEM analysis of SARS-CoV-2-infected tonsil organoids on day 3 in comparison with mock-infected samples. Scale bar, 1,000 nm. (D and E) mRNA expression profile for early (D) and late (E) response genes encoding antiviral proteins, interferons/their receptors, and cytokines or chemokines in SARS-CoV-2-infected tonsil organoids. Colored bar represents fold changes to mock-infected tonsil organoids.

To analyze changes in gene expression induced by SARS-CoV-2 infection in tonsil organoids, we performed mRNA array analysis using a quantitative multiplex panel composed of cytokine- and interferon response-related genes. We found that among the 252 genes tested in the array, 79 genes were upregulated, whereas 48 genes were downregulated (fold change > |2|). Particularly, transcription of the genes encoding OAS1, IL-6, CXCL8, FOS, IL21R and BMP6 was rapidly enhanced by more than 10 folds within 3 h post-infection. Most of the genes selected here have been reported previously to be over-produced uncontrollably in SARS-CoV-2-infected patients as a consequence of “cytokine storm”. It is noteworthy that OAS1, involved in the innate immune response to viral infection for RNA degradation, is prominently stimulated by SARS-CoV-2 throughout the infection period for 72 h. These omics data stressed that the tonsil organoids model reconstitutes highly orchestrated antiviral defense system by restoring SARS-CoV-2 infection-associated host gene expression (Figure 4D,E and S5A-C).

In summary, human tonsil organoids fully recapitulate the important features of *in vivo* tonsil organs; most notably, SARS-CoV-2 can infect tonsil organoids and replicate vigorously, after which viral particles are secreted, indicating that the entire life cycle of SARS-CoV-2 is discernible using tonsil organoids. Most of all, the therapeutic efficacy of remdesvir was validated in the tonsil organoid model. It demonstrated that the organoids can be used as a preclinical drug screening tool for developing SARS-CoV-2 therapeutics. To the best of our knowledge, this is the first paper to show establishment of tonsil organoids and their feasibility of a respiratory viral infection, representatively SARS-CoV-2. From a clinical perspective, tonsils are an easily accessible intraoral organ for harvesting specimens for SARS-CoV-2 studies without great inconvenience to the patient, unlike kidney, liver, bronchus, or gut samples which require invasive and aggressive procedures to obtain. Thus, tonsil organoids may be a valuable *ex vivo* model for SARS-CoV-2 research.

## Discussion

The oropharynx, being located behind the oral cavity and beneath the nasopharynx, functions in both the digestive and respiratory systems. Among the four subparts of the oropharynx, the tonsils (named the palatine tonsils or the oropharyngeal tonsils) are composed of a large population of immune cells and thus responsible for the front line of defense against bacteria and viruses that invade the nasal or the oral cavity. Paradoxically, they also serve as a reservoir for infection of diverse viruses including Epstein-Barr virus, adenoviruses, influenza A and B viruses, herpes simplex virus, rhinovirus, enterovirus and even human papillomavirus (Brook, 1987; Chatterjee et al., 2019; Mellin et al., 2002). Recently, it has been reported that the two major genes required for SARS-CoV-2 entry, ACE2 and TMPRSS2, are expressed in the squamous epithelium lining oropharyngeal tonsillar tissue (Hou et al., 2020). It motivated us to explore if the tonsil tissue is susceptible to the SARS-CoV-2 infection. However, the lack of culture system of tonsillar epithelial cells hindered in-depth cytopathologic or virological investigation with the ongoing pandemic virus. Fortunately, we accomplished establishment of a reliable culture system of the human tonsil organoids, recapitulating the dominant features observed in *in vivo* tonsil organs, such as stratified squamous epithelial layers, tonsillar biomarker distributions, and reproducible gene expression (Figure 2). In addition, differentiation from ITGA6^+^ cells identified their potential origin to be tissue-resident stem cells. Most notably, the tonsil organoids, of which basal layer was occupied with ACE2 and TMPRSS2 proteins, were efficiently infected with SARS-CoV-2. Consequently, it was verified that this virus is vigorously amplified there through multi-round infections, resulting in secretion of the progeny viral particles abundantly (Figure 3 and 4). Being supportive to a well-known theory highlighting the key roles of an oral-lung aspiration axis for airways infectious diseases, our finding addresses that the tonsils might be not only a putative reservoir but also one of the major target organs for primary infection of SARS-CoV-2 before its diffusion to distal intrapulmonary regions.

In actual clinical practice, the specimens are generally collected either by following the nasal or oral route for the COVID-19 test, undesirably resulting in a false-negative rate of 20-67%(Kucirka et al., 2020; Wiersinga et al., 2020). One of the methods to improve the diagnostic accuracy could be to retrieve adequate amounts of viral samples from the correct and accessible site where viral replication occurs actively. However, nasal sample collection from the nasopharynx mucosa is not easy, as it is located deep inside the nasal cavity. A thrust with a cotton swab could make the patients feel uncomfortable. Due to the unexposed nature of the nasopharynx, collecting mucosa from the nasal cavity but not from the nasopharynx could unwantedly happen. Even if viral particles reach the nasal mucosa, functional ciliary movement of the epithelial surface promotes mucus flow physically clearing away the viral contaminants from the nasal mucosa to the nasopharynx. Besides as seen our data with the nasal cavity mucosa organoids, viral replication occurs poorly within the nasal cavity compared to the tonsils. Therefore, it is recommended that specimen collection is performed from the tonsils preferentially, as they are located at a relatively conspicuous site and adjustable for prolonged SARS-CoV-2 replication. More importantly, this approach could alleviate the pain of the patients.

Cytokine storm has been regarded as an initial cause for acute respiratory distress syndrome (ARDS) and multi organ failures accounts for severe pneumonia, accounting for a significant number of deaths among SARS-CoV-2-infected patients. This hyperactive inflammatory response is mainly related to abrupt release of pro-inflammatory or inflammatory mediators such as IL-6, TNF-α and CXCL8 (also named IL-8) in COVID-19 patients (Huang et al., 2020). Beyond this observation, our gene expression profile in tonsil organoids additively signified the elevated mRNA levels of FOS, IL-21R, BMP6, CSF2 and CXCL6 at an earlier point post-infection, together with IFNAR1 and CXCL1 at a later point after SARS-CoV-2 infection. Intriguingly, in accordance with published data from nasal goblet and ciliated cells in the patients (Sungnak et al., 2020), expression of OAS1 mRNA was remarkably stimulated during the whole infection period, indicating operation of aggressive tonsillar innate immune responses for non-specific degradation of both viral and cellular RNAs. On the basis of the immune network information acquired, we are going to investigate the time-course of single-cell RNA-seq. transcriptome changes and the cytokines/chemokines secretion by co-culturing the tonsil organoids in the presence of tonsillar lymphocytes to elucidate immunological tripartite crosstalk between tonsil epithelial cells, lymphocytes and SARS-CoV-2. This approach could provide scientific insights or knowledge for understanding of host defense mechanism against the virus or of counteracting viral immune evasion strategies.

It should be emphasized that the therapeutic efficacy of remdesvir was validated in the tonsil organoid model, implying its value as a preclinical drug screening tool for development of SARS-CoV-2 therapeutics. This system might have advantages as an alternative to animal models particularly when they target host factors involved in the virus life cycle or immune regulations. It is nontrivial because efficacy of those antivirals could be devaluated or not assessable in animals due to low genetic homology or their null function. To the best of our knowledge, this is the first paper to show generation of human tonsil organoids and their feasibility of a respiratory viral infection, representatively SARS-CoV-2. From a clinical perspective, tonsils are an easily accessible intraoral organ for harvesting specimens for SARS-CoV-2 studies without great inconvenience to the patient, unlike kidney, liver, bronchus, or gut samples which require invasive and aggressive procedures to obtain. Thus, tonsil organoids may be an attractive *ex vivo* model for fundamental or applied research on SARS-CoV-2 infections or other microbial infectious diseases.

## Acknowledgements

This work was supported by a grant of the Korea Health Technology R&D Project through the Korea Health Industry Development Institute, funded by the Ministry of Health & Welfare, Republic of Korea (HR16C0002, HI18C2458 to J.Y.), by the Basic Science Research Program through the National Research Foundation of Korea (NRF) funded by the Ministry of Science & ICT, Republic of Korea (2018R1D1A1A02050030 to J.Y., 2016R1A5A2012284 and 2018R1D1A1A02086084 to Y.C.L.), by the 3D-TissueChip Based Drug Discovery Platform Program through the Korea Evaluation Institute of Industrial Technology funded by the Ministry of Commerce, Industry and Energy (20009773 to J.Y.), by a National Research Foundation of Korea (NRF) grant funded by the Korean government (MSIT) (NRF-2020M3A9I2081687 to M.K.) and by an intramural research funding from KRICT (KK2032-20 to M.K.). The SARS-CoV-2 resource (NCCP No., 43326) was provided by the National Culture Collection for Pathogens, Republic of Korea.

## Author Contributions

M.K., Y.C.L. and J.Y. designed and supervised the study. H.K.K., H.K., W.H.C. and J.-Y.P. performed the experiments. H.K.K, H.K. and W.H.C. conducted the histological staining. K.H.K. and H.W.H. analyzed the genomics data. M.K.L, Y.J. and J.S.S performed SARS-CoV-2-related experiments in a BSL3 facility. Y.C.L. provided the human tissues and clinical specimens. H.K.K., S.I.H., M.K., Y.C.L. and J.Y. wrote the manuscript.

## Declaration of Interests

The authors declare no competing interests.

## Supplementary Materials

### Materials and Methods

#### Cell isolation and organoid formation from human tonsil tissue

This study was approved by the institutional review board of Konkuk University Hospital (IRB-No.KUH1110073) and carried out with the written consent of all donors. Tonsils were obtained from patients by tonsillectomy. When needed, both tonsil and nasal tissues were collected from one patient *via* tonsillectomy and endoscopic nasal surgery, respectively, for the co-treatments of tonsillitis and sinusitis. The samples were chopped and washed with D-PBS (LB001-02, Welgene, Daegu, Korea) and then enzymatically digested with advanced DMEM/F12 (11330-032, Gibco, Grand Island, NY, USA) containing 1 mg/mL collagenase II (17101015, Gibco) for 2 h at 37°C. After digestion, isolated cells were embedded in Matrigel (354230, Corning, Inc., Corning, NY, USA), seeded into a 48-well plate (SPL, Inc., Gyeonggi-do, Korea) and incubated with 5% CO_2_ at 37°C for 10 min to polymerize the matrices. Tonsil organoids were cultured in advanced DMEM/F12 supplemented with Antibiotic-Antimycotic (Thermo Fisher Scientific, Fisher Scientific, Waltham, MA, USA), Glutamax (Thermo Fisher Scientific), B27 (Invitrogen, Carlsbad, CA, USA), 10% R-spondin1 conditioned media and the following growth factors: 50 ng/mL recombinant murine HGF (315-23, Peprotech, Rocky Hill, NJ, USA), 100 ng/mL noggin (cyt-600, ProSpec, St. Paul, MN, USA), 20 nM A83-01 (SML0788, Sigma, St. Louis, MO, USA), 50 ng/mL human FGF10 (ATGP1387, ATGen, Seongnam, Korea), 20 ng/mL human bFGF (100-18B, Peprotech), 10 µM prostaglandin E2 (3632464, BioGems, Westlake Village, CA, USA), and 10 mM nicotinamide (N0636, Sigma). Five nM Neuregulin1 (100-03, Peprotech) was added only to the nasal cavity mucosa derived organoid tonsil cultures. After passaging, 10 µM Y-27632 (1254, Tocris Biosciences, Bristol, UK) was added to the culture medium for 2 days.

#### Infection of tonsil organoids with SARS-CoV-2

SARS-CoV-2 (BetaCoV/Korea/KCDC03/2020) was amplified in Vero CCL-81 cells (American Type Culture Collection, Rockville, MD, USA) through three passages in Dulbecco’s modified Eagle’s medium (DMEM; HyClone, South Logan, UT, USA). Tonsil organoids at passage 2 were differentiated from single cells for 5 days. After detaching the organoids with 6 × 10^4^ cells, they were suspended in 5 μL TeM and infected with the same volume of SARS-CoV-2 at an MOI of 0.1 or 1 at 37°C for 1 h. Unabsorbed virus was removed by washing with 1 mL PBS twice after centrifugation at 1,000 *g* for 10 s. Each sample of mock-infected or SARS-CoV-2-infected organoids was embedded into 20 μL Matrigel on a 48-well plate and cultured in 300 μL TeM supplemented with 10 μM Y27632 at 37°C. The supernatants were harvested at 3, 24, 48, and 72 h post-infection for viral RNA quantification and infectious viral titration.

#### Viral RNA quantification and TCID_50_ determination

Viral RNA was purified from the culture supernatant at different time points using a Viral RNA Purification Kit (Qiagen, Hilden, Germany). One-step RT-PCR targeting the viral N gene was performed using a diagnostic kit (PCL Inc., Seoul, Republic of Korea) and CFX96 Touch real-time PCR instrument (Bio-Rad, Hercules, CA). In all experimental sets, RNAs purified from serial dilutions of a virus stock with identified plaque titers were used as a standard to quantify the absolute viral RNA copy number in the samples. In parallel, Vero CCL-81 cells (ATCC) were seeded at a density of 2 × 10^4^ cells per well in 96-well plates, and then treated with serially diluted culture supernatants from the SARS-CoV-2-infected organoids using the cell-cultured SARS-CoV-2 stock at an MOI of 0.1 as a control. On day 2 post-infection, the cells were fixed and permeabilized with chilled acetone: methanol (1:3) solution at room temperature for 10 min. Viral S protein was probed using anti-S antibody (Genetex, Irvine, CA, USA) and Alexa Fluor 488-conjugated goat anti-mouse IgG antibody (Invitrogen), and cellular nuclei were stained with 4’,6-diamidino-2-phenylindole (Invitrogen). Fluorescence images were captured and analyzed using an Operetta High-content Screening System (PerkinElmer, Waltham, MA, USA) and quantified using the built-in Harmony software. Compared to the control infection (100%), dilution folds resulting in 50% infection were calculated to determine the 50% tissue culture infectious doses (TCID_50_).

#### Antiviral activity of remdesivir against SARS-CoV-2 in tonsil organoids

Five-day cultured tonsil organoids were infected with SARS-CoV-2 at an MOI of 0.1 and embedded in 20 μL fresh TeM plus Matrigel (6 × 10^4^ cells per well in 48-well plates) after washing twice with PBS. Remdesivir (purity 99.84%; MedChemExpress, Monmouth Junction, NJ, USA) was 10-fold serially diluted in TeM from 10 to 0.1 μM to treat SARS-CoV-2-infected organoids. On day 2, the culture supernatants were harvested to measure the amount of viral RNA, in parallel cell lysates were collected from 4 wells to compare the expression level of viral S protein. For western blot analysis, cell lysates were harvested using RIPA buffer (LPS solution, Daejeon, Republic of Korea). The immuno-transferred membrane was probed with primary anti-S antibody followed by secondary horseradish peroxidase-conjugated anti-mouse goat IgG (Invitrogen).

#### Clonal organoid formation assay from sorted cells by flow cytometry

To confirm the origin of cells forming the organoids, tonsil tissue was enzymatically digested with advanced DMEM/F12 medium containing 600 µg/ml collagenase II and 250 µg/ml DNase I (D5025, Sigma-Aldrich) for 2 hours at 37°C. Dissociated cells were stained with anti-IgG (130-113-196, Miltenyi Biotec), NGFR (sc-13577, Santa Cruz), CD44-FITC (555478, BD biosciences), ITGA6-PE (12-0495-83, Biolegend), and EpCAM-APC (130-113-260, Miltenyi Biotec), Alexa 488 (A32723, Invitrogen), and sorted using the MACS method. Sorted cells were embedded in Matrigel and cultured in TeM (Full name) for 15 days.

#### Chemotaxis assay and cytokine array

To demonstrate the chemoattracts production of tonsil organoids, the chemotaxis assay was performed. Briefly, HL-60 cells (10240, Korean Cell Line Bank) were differentiated into granulocytes by adding 1.3% DMSO (D1370.0250, Duchefa) for 7 days, as previously described^49^. And then, HK-60 cells monolayers grown on transwell membranes (3421, Corning) at a density of 5×10^5^ cells/well in RPMI 1640 (11875093, Gibco) medium containing 0.5% BSA, were exposed to supernatants that were collected from LPS (10 µg/ml, L2630, Sigma) treated organoids. After 1 hour, migrated HL-60 cells were counted using LUNA™Automated Cell Counter (Biosystems). The cytokine profile assay was performed using a Proteome Profiler Human Cytokine Array kit (ARY005B, R&D Systems). Briefly, culture supernatants from LPS-treated and untreated tonsil organoids was collected and incubated with the precoated Proteome Profiler array membrane at 2-8°C overnight. The array membrane was washed and incubated with streptavidin-HRP buffer for 30 min. And then, membrane was treated with Chemi Reagent Mix and dot densities were imaged on a LAS4000.

#### Microarray analysis

To analyze the genetic similarity between tonsil tissue and organoids, microarray was performed according to the human microarray chip technical manual (Macrogen, Seoul). Briefly, total RNA was extracted from tissue and organoids using MagListo™ 5M Cell Total RNA Extraction Kit (K-3611, Bioneer) according to the manufacturer’s protocol. Quality control and gene expression profiling were performed at Macrogen. Gene expression data were processed using Affymetrix Power Tools and R statistical software version 3.2.2.

#### qRT-PCR analysis

Total RNA was extracted from LPS-treated or SARS-CoV-2 infected organoids, and cDNA synthesis was conducted using PrimeScript™ RT Master Mix (RR036A, TaKaRa). Quantitative RT-PCR was performed with a Thermal Cycler Dice® Real-Time System III (Takara) using SYBR® Premix Ex Taq™ II (RR820A, TaKaRa). The specific primer sets for Antiviral Response (SH-0014), Chemokines and Receptors (SH-0035) and Interferons and Receptors (SH-0092), which were purchased from Bioneer (AccuTarget™ Human qPCR Screening Kit, Korea). Other primer sequences are listed in Table S2.

#### Histology, immunohistochemistry, and immunofluorescence

Tissues and organoids were fixed in 4% paraformaldehyde (PFA, BP031, Bio-solution), and embedded in paraffin. Paraffin sections of 5 µm in thickness were deparaffinized in xylene and hydrated in a graded series of ethanol. These samples were then stained with H&E, Alcian blue (AR16011-2, Agilent), PAS staining kit (ab150680, Abcam), and Masson’s trichrome staining kit (QAR17392-2, Dako) according to their respective manufacturer’s protocol. For immunohistochemistry, antigen retrieval was performed using sodium citrate buffer (10 mM sodium citrate with 0.05% Tween 20, pH 6) at 95°C for 1 hour. Endogenous peroxidase was blocked using 3% H_2_O_2_ in methanol for 10 min. After blocking with 1% BSA in PBS, sections were incubated at 4°C overnight with primary antibody and then incubated with a biotinylated secondary antibody (anti-mouse IgG or anti-rabbit IgG, Vectastain ABC kit, Vector) at room temperature (RT) for 30 min. Sections were stained according to the avidin-biotin complex method using Vectastain kit and visualized using 3,3′-diaminobenzidine (DAB) (Vectastain DAB kit, Vector). For immunofluorescence analysis, fixed samples were cryoprotected by immersing in PBS containing 30% sucrose and 0.1% sodium azide at 4°C, embedded in optimal cutting temperature (OCT, 4538, Sakura) compound, and rapidly frozen in liquid nitrogen. Frozen sections of 4 µm in thickness were pre-blocked with 5% normal horse serum (S-2000, Vector) in PBS at RT for 2 hours and incubated with primary antibody at 4°C overnight. And then, sections were incubated with secondary antibody for 2 hours at RT. Nuclei were counterstained with Hoechst 33342 (1 µg/ml, B2261, Sigma). The acquisition of confocal images was performed by Zeiss LSM 880 confocal microscopy. All antibodies used in this study are listed in Table S3.

#### Transmission electron microscope (TEM)

To observe the SARS-CoV-2 infection into tonsil organoid, transmission electron microscope (TEM) analysis was performed. Briefly, tonsil organoids were fixed in Karnovsky’s fixative (2% glutaraldehyde, 2% paraformaldehyde, 0.5% CaCl_2_ in 0.1 M phosphate buffer, pH 7.4), and embedded in the epoxy resin after post-fixation with 1% OsO4 dissolved in 0.1 M phosphate buffer. Ultrathin sections were processed using an ultramicrotome (EM-UC-7, LEICA) and placed on a copper grid stained with 6% uranyl and lead citrate. The samples were observed using a JEM-1011 (JEOL).

#### Statistical analysis

Statistically significant differences were analyzed using GraphPad Prism software package, version 3.0 (GraphPad Prism). Unpaired two-tailed t-tests were used to compare two groups and one-way ANOVA with post-test Tukey multiple comparisons tests were used to compare multiple groups. Significance was considered at p < 0.05. Sample sizes for experiments were estimated using NIS imaging software. All results are presented as the mean ± SEM.

**Table S1.** Medium composition for organoid establishment.

| Reagent | HCM | HPM | TfM | TeM | NeM |
| --- | --- | --- | --- | --- | --- |
| Advanced DMEM/F12<br>(Thermofisher, 12634-010) | Basal<br>Medium | Basal<br>Medium | Basal<br>Medium | Basal<br>Medium | Basal<br>Medium |
| B27 Supplement<br>(Thermofisher, 17504044) | 1X | 1X | 1X | 1X | 1X |
| Glutamax<br>(Thermofisher, 35050-061) | 1X | 1X | 1X | 1X | 1X |
| Antibiotics/Antimycotics<br>(Welgene, LS203-01) | 1X | 1X | 1X | 1X | 1X |
| R-spondin1 conditioned medium<br>(Homemade) | 10% | 10% | 10% | 10% | 10% |
| Human FGF10<br>(ATGen, ATGP1387) | 50 ng/ml | 50 ng/ml | 50 ng/ml | 50 ng/ml | 50 ng/ml |
| Human Gastrin<br>(Sigma, G9145) | 50 ng/ml |  |  |  |  |
| HEPES<br>(Thermofisher, 15630-080) | 1mM |  |  |  |  |
| A83-01<br>(Sigma, SML0788) | 20nM | 20nM | 20nM | 20nM | 20nM |
| Wnt3a conditioned medium<br>(Homemade) | 10% |  |  |  |  |
| Nicotinamide<br>(Sigma, 72340) | 10 mM | 10 mM | 10 mM | 10 mM | 10 mM |
| Human Noggin<br>(Prospec, cyt-600) | 100 ng/ml | 100 ng/ml | 100 ng/ml | 100 ng/ml | 100 ng/ml |
| Human EGF<br>(Peprotech, AF-100-15) | 50 ng/ml | 50 ng/ml |  |  |  |
| Human FGF2<br>(Peprotech, 100-18B) |  | 20 ng/ml | 20 ng/ml | 20 ng/ml | 20 ng/ml |
| Human HGF<br>(Peprotech, 315-23) |  |  |  | 50 µg/ml | 50 µg/ml |
| Human Neuregulin-1<br>(Peprotech, 100-03) |  |  |  |  | 5nM |
| SB202190<br>(Sigma, s7076) |  | 10 µM | 10 µM |  |  |
| Prostaglandin E2<br>(BioGems, 3632464) |  | 10 µM | 10 µM | 10 µM | 10 µM |

\*Abbreviations: HCM; Human colon organoid medium, HPM; Human prostate organoid medium, TfM; Tonsil organoid formation medium, TeM; Tonsil organoid expansion medium, NeM; Nasal cavity mucosa-derived organoid expansion medium

**Table S2.**
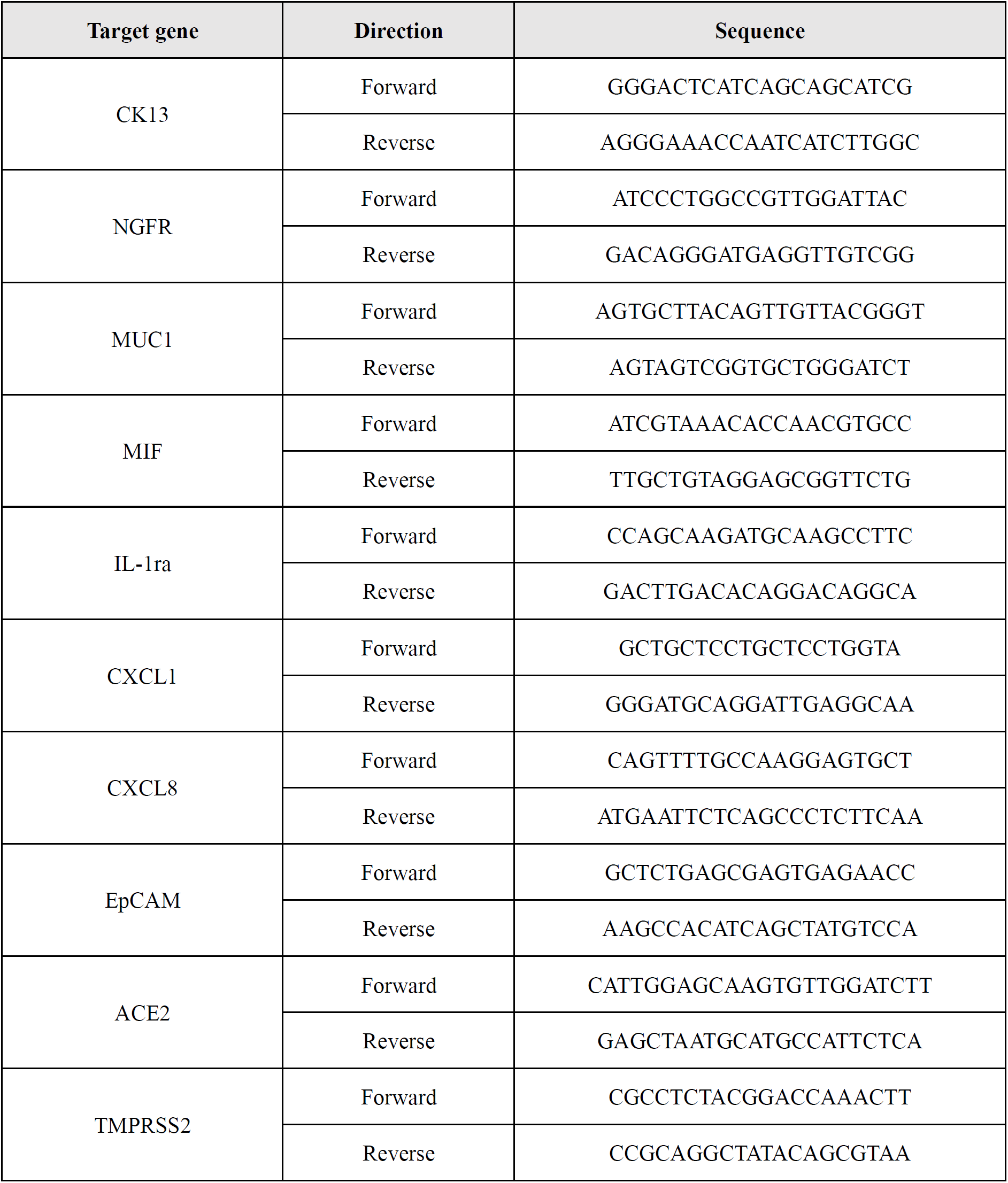
Primers list for qRT-PCR analysis.

**Table S3.** List of antibodies used for immunostaining studies.

| Target Antigen | Host | Catalog No. | Company | Dilution |
| --- | --- | --- | --- | --- |
| Cytokeratin 4 | Mouse | Sc-52321 | Santa Cruz | 1 : 350 |
| Cytokeratin 5 | Mouse | Sc-81702 | Santa Cruz | 1 : 250 |
| NGFR | Mouse | Sc-13577 | Santa Cruz | 1 : 200 |
| CD44 | Mouse | BD550988 | BD biosciences | 1 : 120 |
| CD44 | Mouse | BD555478 | BD biosciences | 1 : 100 |
| ITGA6 | Rabbit | HPA012696 | Sigma | 1 : 400 |
| ITGA6 | Rat | 12-0495-83 | Biolegend | 1 : 100 |
| Ki67 | Rabbit | Ab16667 | Abcam | 1 : 250 |
| Muc1 | Mouse | BD555925 | BD biosciences | 1 : 150 |
| E-cadherin | Rabbit | Sc-7870 | Santa Cruz | 1 : 300 |
| E-cadherin | Goat | AF748 | R&D systems | 1 : 100 |
| EpCAM | Mouse | Sc-59906 | Santa Cruz | 1 : 300 |
| BrdU | Rat | NB500-169 | Novus Biologicals | 1 : 300 |
| ACE2 | Rabbit | GTX101395 | Genetex | 1 : 400 |
| TMPRSS2 | Rabbit | GTX64544 | Genetex | 1 : 200 |
| dsRNA | Mouse | 10010200 | Scicons | 1 : 250 |
| SARS-CoV /<br>SARS-CoV-2 Spike | Mouse | GTX632604 | Genetex | 1 : 500 |

**Figure S1.**
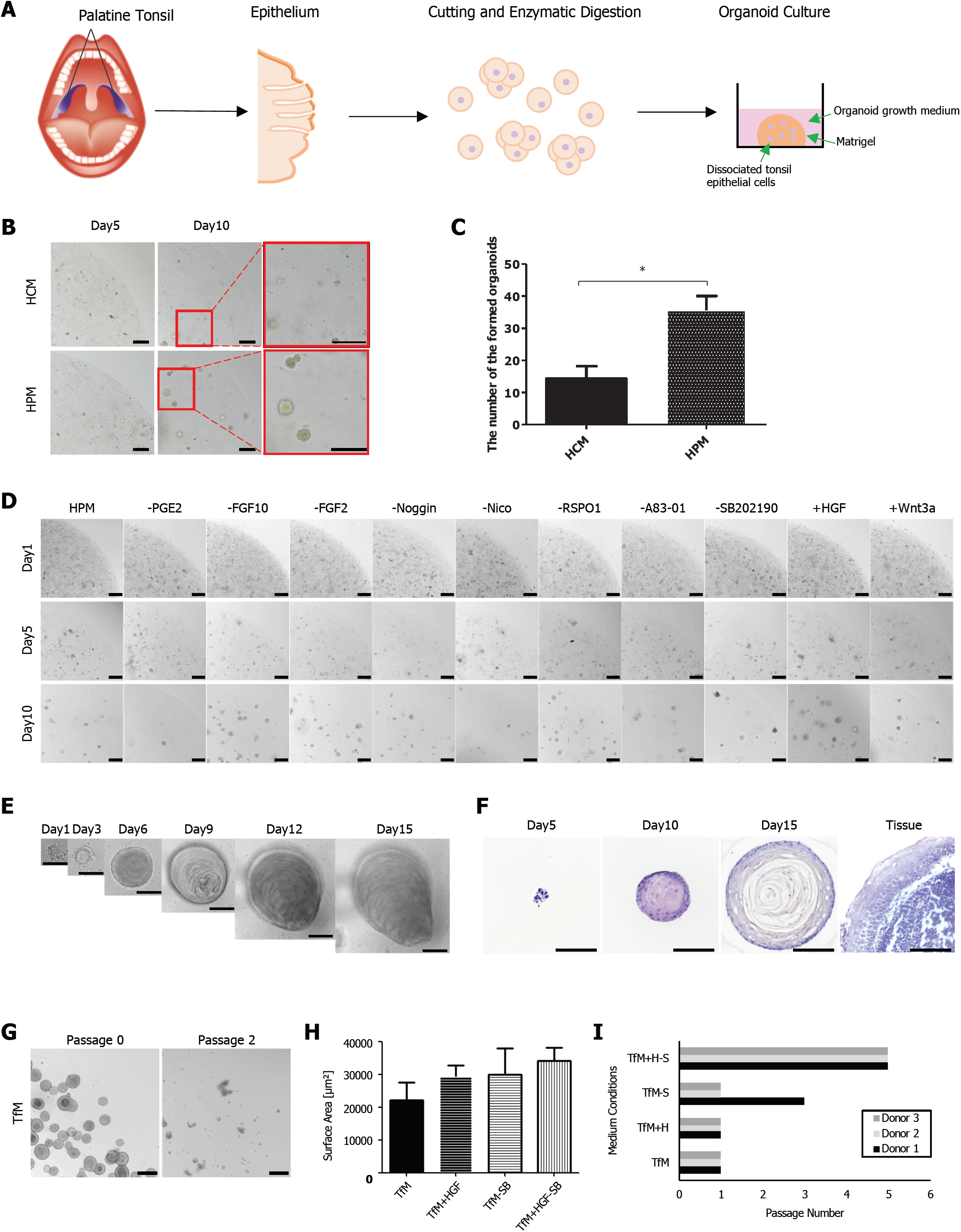
Optimization of a method for establishing human tonsil organoids from tonsil tissue. (A) Schematic showing how the organoid cultures were prepared from human tonsil tissue. (B) Representative images of tonsil organoids cultured *in vitro* with human colon media (HCM) and human prostate media (HPM) on days 5 and 10. (C) The bar graph shows the number of organoids that formed when cells were cultured in HCM or HPM. The data represents the mean ± SEM of triple replicates. (D) Representative images of human tonsil organoids cultured in HPM containing various combinations of the indicated factors. (E) The expansion time frame for organoids cultured in TfM. (F) H&E staining of organoids cultured in TfM and human tonsil tissue. (G) Representative images of passage 0 and 2 of organoids cultured in TfM. (H) The surface area of organoids cultured in TfM, TfM adding hepatocyte growth factor (HGF), TfM removing SB202190, and TfM adding HGF and removing SB202190. (I) Final passage numbers reached by tonsil organoids cultured in four different medium. In all images, the scale bar is 100 µm.

**Figure S2.**
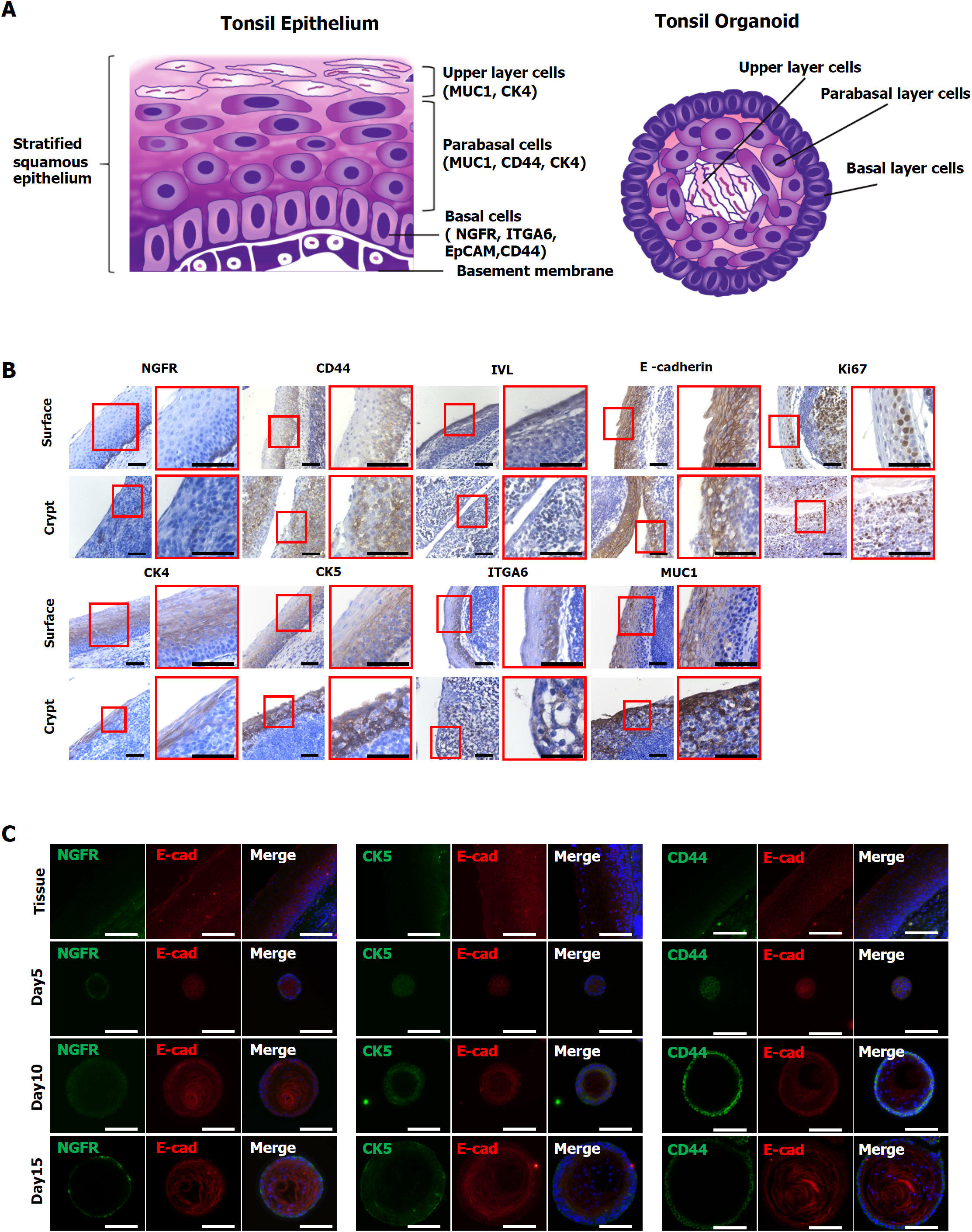
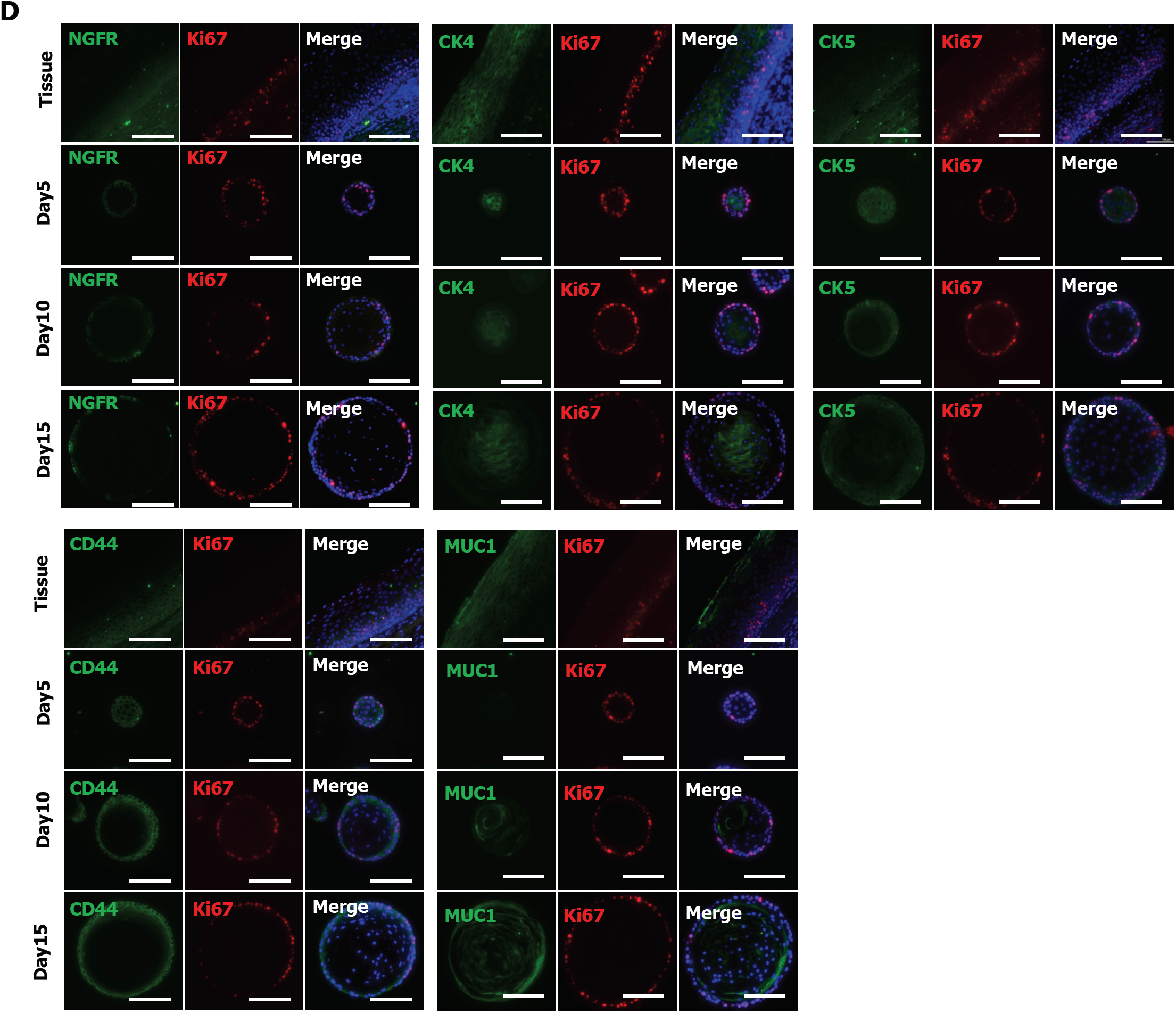
Structural characterization of tonsil organoids. (A) Illustration showing structure and markers for each layer of tonsil epithelium and tonsil organoids. (B) Immunohistochemistry staining of surface and crypt regions in the tonsil epithelium with tonsil epithelium markers (CD44, NGFR, CK4, CK5, E-cadherin, MUC1, ITGA6, IVL) and a proliferation marker (Ki67). Scale bar: 50 µm. (C) Immunofluorescence staining of NGFR, CK5 or CD44 with E-cadherin in human tonsil epithelium and organoids cultured in TeM on days 5, 10, and 15. Scale bar, 100 µm. (D) Immunofluorescence staining of NGFR, CK4, CK5, CD44 or MUC1 with Ki67 in human tonsil epithelium and organoids cultured in TeM on days 5, 10, and 15. Scale bar, 100 µm. *Abbreviations: NGFR; Nerve growth factor receptor, CK4; Cytokeratin 4, CK5; Cytokeratin 5, ITGA6; Intergrin-α6, IVL; Involucrin

**Figure S3.**
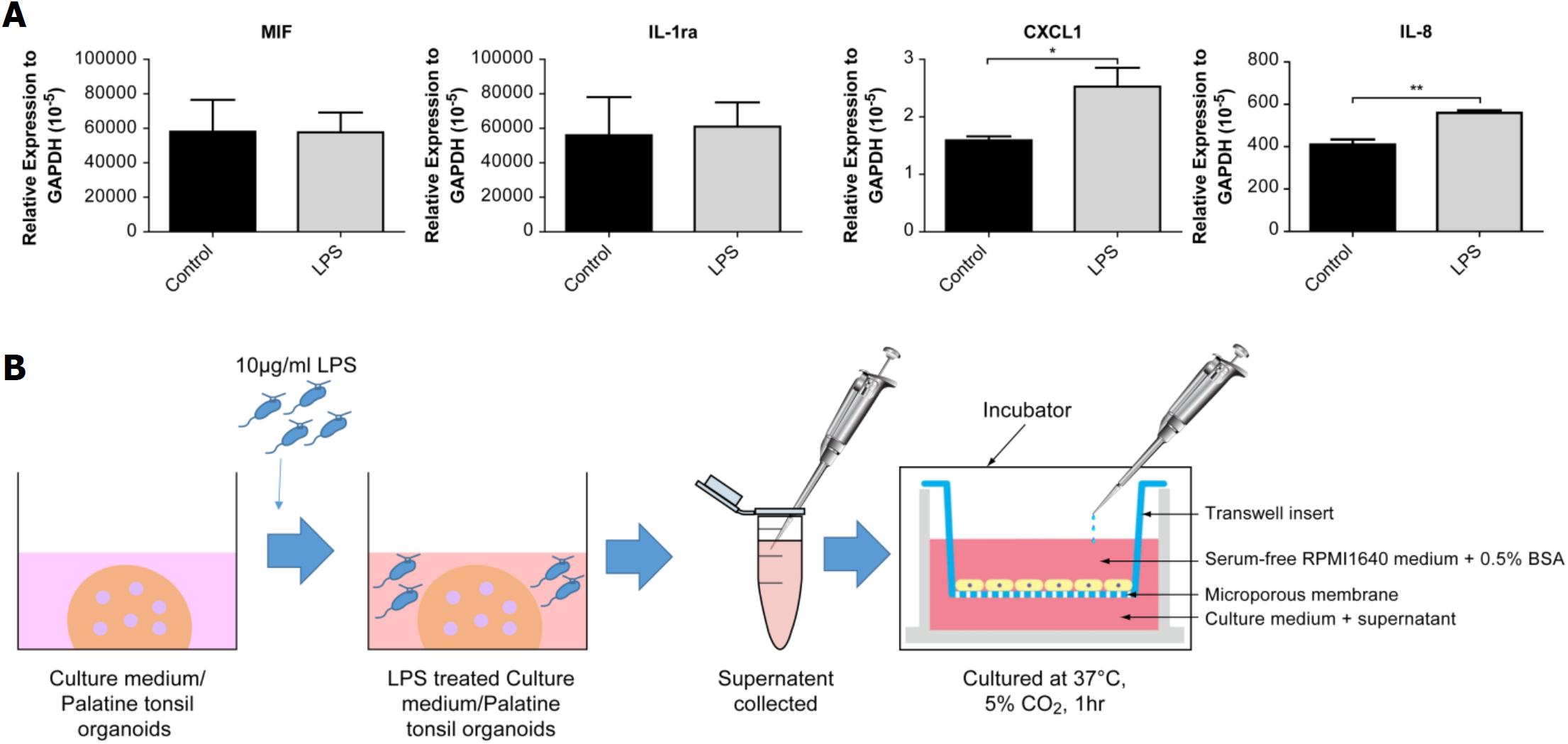
Tonsil organoids release cytokines in response to infection stimuli. (A) qPCR analysis of cytokines (MIF, IL-1ra, CXCL1, IL-8) released from human tonsil organoids stimulated with 10 µg/ml LPS. Data are presented as the mean values of triple replicates ±SEM. **p<0.01, *p<0.05, according to t-test. (B) Illustration showing the migration assay of HL-60 cell line using conditioned media from LPS-stimulated tonsil organoids *Abbreviations: MIF; Macrophage migration inhibitory factor, IL-1ra; Interleukin-1α, IL-8; Interleukin-8

**Figure S4.**
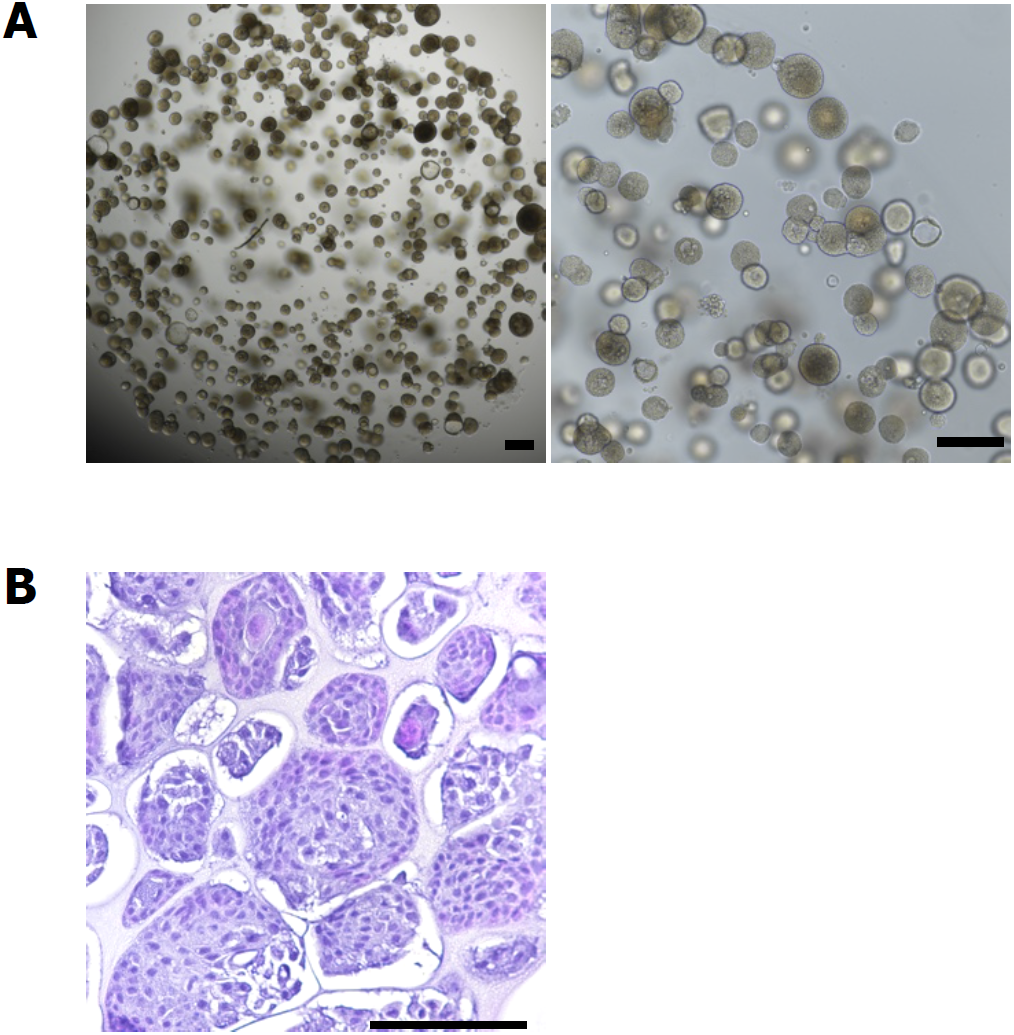
Establishment of human nasal cavity mucosa-derived organoids. (A) Representative images of human nasal cavity epithelium-derived organoids cultured *in vitro* with human nasal cavity mucosa-derived organoid expansion media (NeM). (B) H&E staining of human nasal cavity epithelium-derived organoids cultured in NeM on days 15.

**Figure S5.**
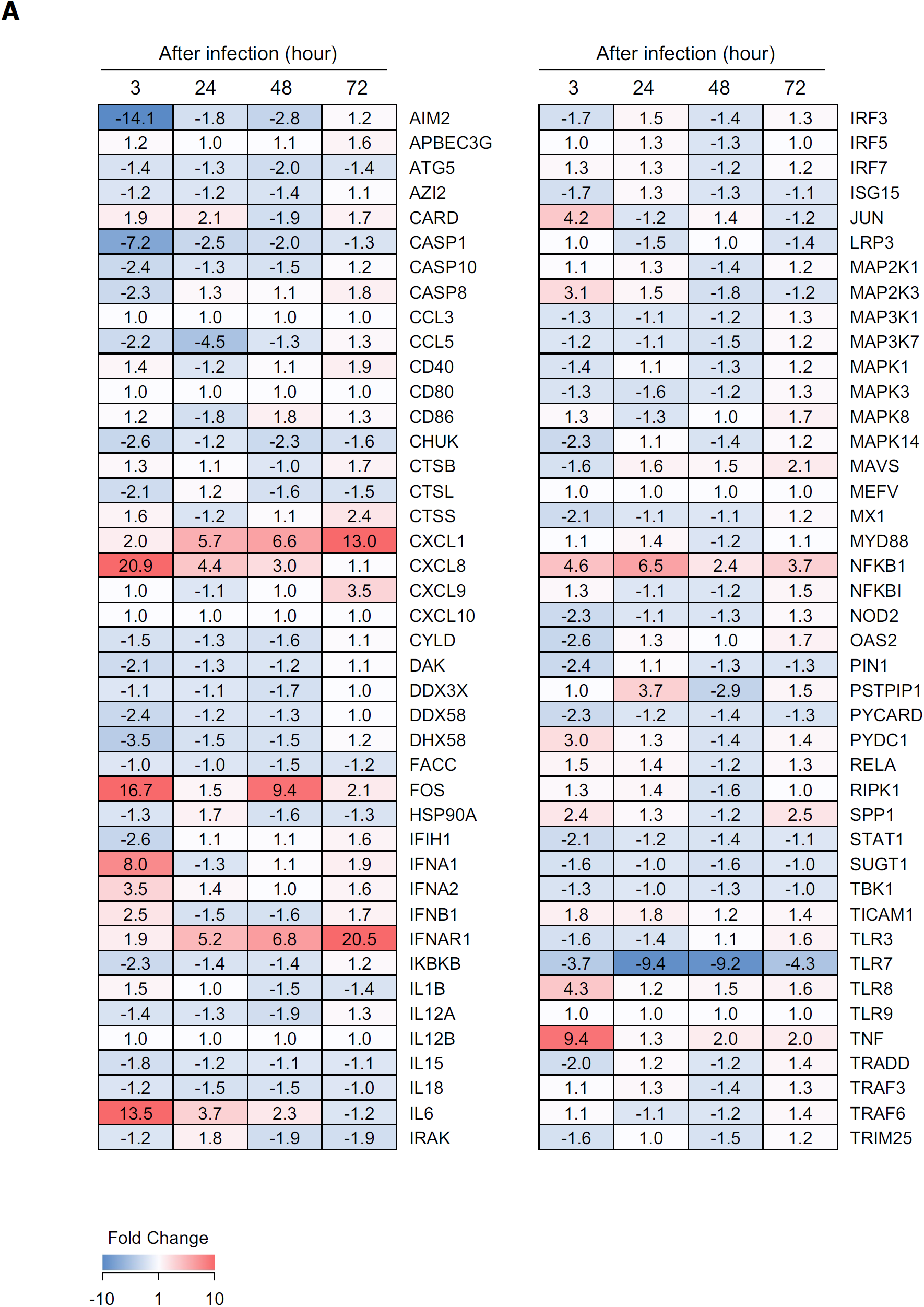
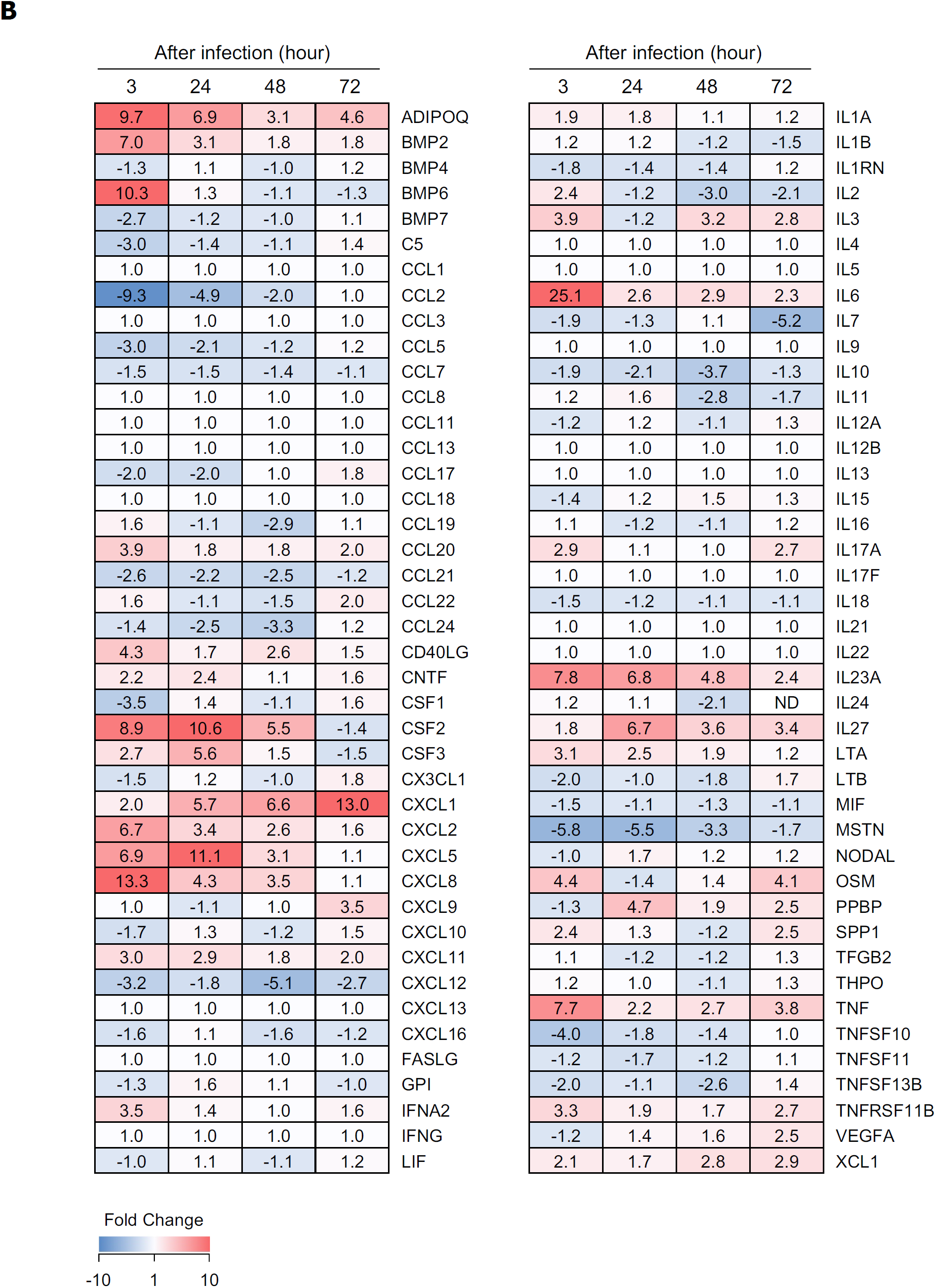
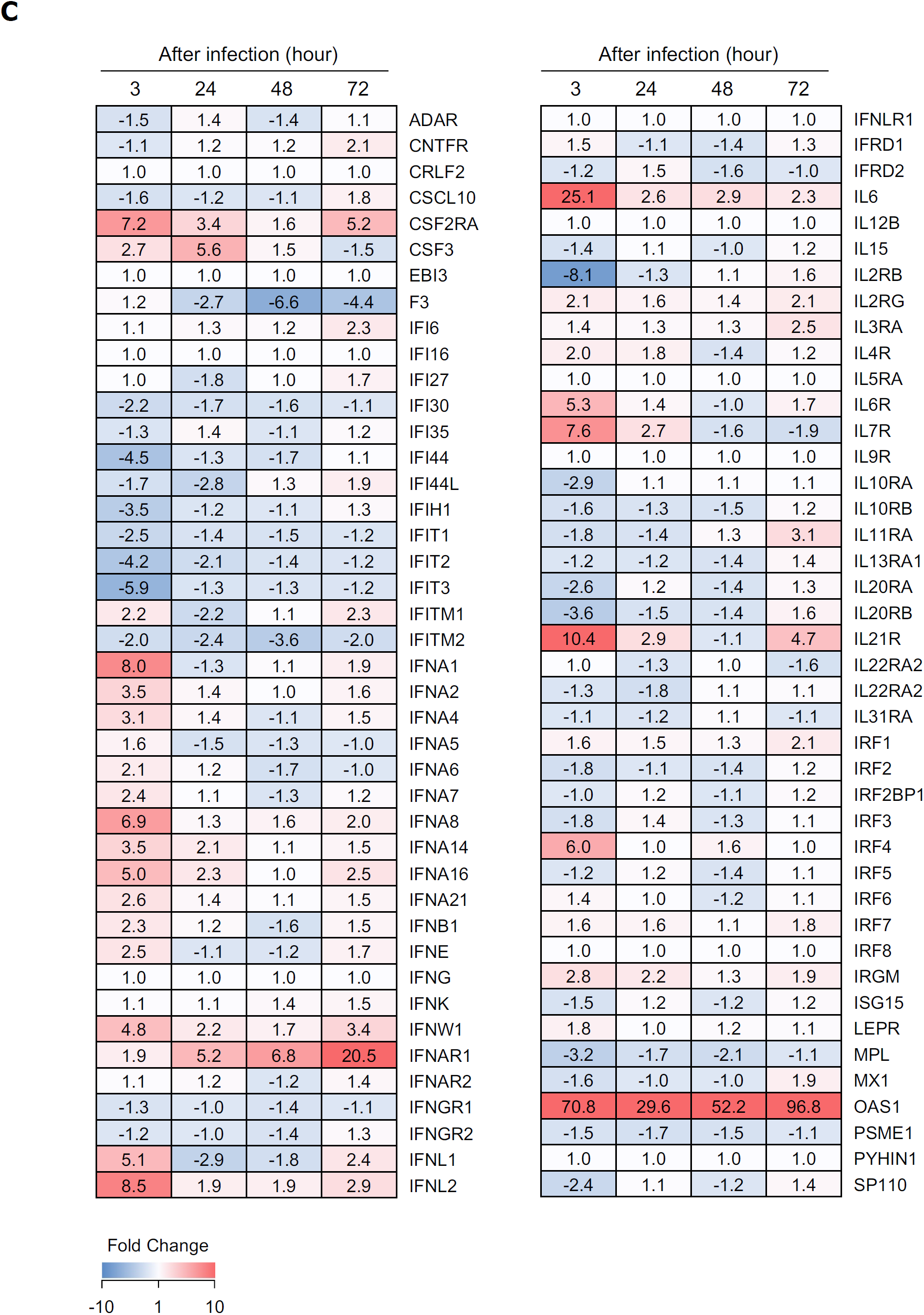
mRNA expression profile in SARS-CoV-2 infected tonsil organoids. Heatmaps depicting fold changes in mRNA expression in SARS-CoV-2 infected tonsil organoids against mRNA expression in uninfected tonsil organoids. (A) The selected genes in this experiment are related to the antiviral responses. Colored bar represents fold changes. (B) The selected genes in this experiment are related to the interferon responses. (C) The selected genes in this experiment are related to the cytokines and chemokines.

## Notes

### Competing Interest Statement

The authors have declared no competing interest.

